# Class switching is differentially regulated in RBC alloimmunization and vaccination

**DOI:** 10.1101/2023.01.11.523608

**Authors:** Anupam Prakash, Jelena Medved, Abhinav Arneja, Conrad Niebuhr, Andria N. Li, Soraya Tarrah, Alexis R. Boscia, Emily D. Burnett, Aanika Singh, Juan E. Salazar, Wenhao Xu, Manjula Santhanakrishnan, Jeanne E. Hendrickson, Chance John Luckey

**Author notes:** Corresponding Author Information: Chance John Luckey, 415 Lane Road, Building MR5, Rm 3320, Charlottesville, VA 22903.

## Abstract

**Background:** Studies of human patients have shown that most anti-RBC alloantibodies are IgG1 or IgG3 subclasses, though it is unclear why transfused RBCs preferentially drive these subclasses over others. Though mouse models allow for the mechanistic exploration of class-switching, previous studies of RBC alloimmunization in mice have focused more on the total IgG response than the relative distribution, abundance, or mechanism of IgG subclass generation. Given this major gap, we compared the IgG subclass distribution generated in response to transfused RBCs relative to protein in alum vaccination, and determined the role of STAT6 in their generation.

**Study Design and Methods:** WT mice were either immunized with Alum/HEL-OVA or transfused with HOD RBCs and levels of anti-HEL IgG subtypes were measured using end-point dilution ELISAs. To study the role of STAT6 in IgG class-switching, we first generated and validated novel STAT6 KO mice using CRISPR/cas9 gene editing. STAT6 KO mice were then transfused with HOD RBCs or immunized with Alum/HEL-OVA, and IgG subclasses were quantified by ELISA.

**Results:** When compared to antibody responses to Alum/HEL-OVA, transfusion of HOD RBCs induced lower levels of IgG1, IgG2b and IgG2c but similar levels of IgG3. Class switching to most IgG subtypes remained largely unaffected in STAT6 deficient mice in response to HOD RBC transfusion, with the one exception being IgG2b. In contrast, STAT6 deficient mice showed altered levels of all IgG subtypes following Alum vaccination.

**Discussion:** Our results show that anti-RBC class-switching occurs via alternate mechanisms when compared to the well-studied immunogen alum vaccination.

## Introduction

Red Blood Cell alloimmunization is a major problem for those patients who require chronic transfusions.^1^ Allogenic anti-RBC IgG antibodies can make it difficult to find compatible blood for many patients. For others, production of anti-RBC IgG antibodies can induce delayed hemolytic transfusion reactions (DHTRs) that remain an all too common cause of patient morbidity and mortality.^2,3^ Despite the clinical importance of anti-RBC alloantibodies, we do not completely understand how transfusion of foreign RBCs leads to the production of class-switched IgG antibodies.

IgG antibodies are not homogenous, as individual antibodies are expressed as a particular IgG isotype that has unique effector properties. Expression of different IgG isotypes occurs when B cells are induced to undergo class-switching, a process by which a fixed antigen binding region of the heavy chain locus is rearranged so that it is expressed along with one of four different unique IgG constant regions.^4^ Since each of these constant regions interacts with different effector molecules, class-switching serves as a mechanism to diversify the potential effector functions of a given antibody specificity. Indeed, each IgG isotype has its own unique half-life, serum abundance, affinity for specific Fc receptors, and ability to activate complement. Thus, knowing which isotypes of antibodies predominate in a given immune response can inform you of the potential functional outcome.

Given their functional differences, it is not surprising that the production of specific IgG subclasses is carefully regulated. Though most immune responses generate antibody class switching to all IgG subclasses, the relative amount of each subclass has been shown to differ as a result of specific immunogenic stimuli.^5,6^ In mouse models, viral infections and mRNA vaccinations predominantly drive class switching to IgG2 subtypes in response to Th1 cytokines such as IFN-γ.^7–13^ Parasitic infections and alum vaccination preferentially drive class-switching to IgG1 in response to Th2 cytokines such as IL-4 and IL-13.^12–17^ Encapsulated bacteria and pneumococcal vaccines preferentially favor the IgG3 isotype via a largely T cell-independent mechanism.^18–22^ Thus, IgG isotype distributions can provide important information on the specific class of immune stimuli and resultant cytokine signaling pathways that drive a given antibody response.

There are a remarkably limited number of studies that have looked at the relative amounts of IgG subclasses that are generated in patients in response to RBC transfusion, most dating back to the 1960s and 1970s. The majority of these studies focused on either pregnancy related anti-RhD antibodies or RhD negative volunteers who were intentionally immunized and boosted with RhD positive blood.^23^ There were however a few studies that looked at patient responses to transfusions.^24–26^ Collectively, the predominant isotypes of anti-RBC alloantibodies generated in response to transfusion are IgG3 and IgG1, while IgG2 and IgG4 were either not detected or present at very low titers in the majority of cases.^24–26^ Interestingly, the observed profile of IgG isotypes identified in patients does not fit well with any clear Th1, Th2 or T-independent profile since in humans; where Th1 favors IgG3 and suppresses IgG1, Th2 favors IgG1 and IgG4 while suppressing IgG3, and T independent responses favor IgG2.^27^ Thus, it is unclear how selective IgG isotypes are driven in response to transfusion in patients.

In order to better understand the molecular drivers of RBC alloimmunization and the induction of class-switching to IgG, we turned to the experimentally tractable HOD mouse model of RBC alloimmunization. The HOD model is a widely used system to understand mechanisms of storage-dependent RBC alloimmunization. HOD transgenic mice express a fusion protein consisting of Hen Egg Lysozyme (HEL), Ovalbumin (OVA) and human Duffy on the surface of RBCs.^28^ Multiple studies have shown that transfusion of HOD RBCs into recipient mice after storage leads to a class switched anti-HEL IgG response.^29–32^ Furthermore, we have previously shown that like most infectious stimuli,^5^ HOD transfusion into C57BL/6 mice induces an anti-HEL IgG response consisting of all four of the C57BL/6 mouse isotypes: IgG1, IgG2b, IgG2c and IgG3.^32^ However, our understanding of the molecular regulators of transfusion-induced class-switching remains limited.

To measure the relative ability of transfused RBCs to induce a given IgG subtype, we directly compared HOD RBC transfusion with HEL-OVA protein emulsified in Alum. Protein in Alum vaccination is a well-studied immunization model that has served as the gold standard for the study of IgG class switching in mice. This allowed us to compare and contrast the class-switching observed in response to transfusion with that observed in response to Alum vaccination.

To better understand the molecular mechanisms regulating IgG isotype specific class switching in response to RBC transfusion, we further tested the impact of STAT6 deficiency on class switching to both transfusion and Alum vaccination. STAT6 is a well-known regulator of class switching, where, in mice it supports IgG1 and suppresses IgG2 subtypes.^14,33–45^ Thus, transfusion of STAT6 KO mice allows us to directly test whether the molecular mechanisms that regulate class-switching in response to RBC transfusion are similar or different from those that regulate class-switching in response to protein in Alum vaccination.

## Materials and Methods

### Mice

8-10 weeks old C57BL/6J mice (Jackson Laboratories Strain # 000664) and STAT6 KO mice (described below) were housed at the University of Virginia Animal Care Facility. HOD transgenic mice were maintained on an FVB background as previously described.^28^ HOD transgenic RBCs contain the triple fusion protein of Hen Egg Lysozyme, Ovalbumin and Duffy. All mouse protocols were approved by Institutional Animal Care and Use Committees of University of Virginia, Charlottesville.

### Generation of STAT6 KO mice

CRISPR/cas9 genome editing technology was used to generate STAT6 KO mice. Three gRNAs targeting SH2 coding exons were selected using Desktop Genetics (DESKGEN) guide picker.^46^ crRNA, tracrRNA and Cas9 were purchased from IDT (Coralville, Iowa). The crRNA sequences are listed here: crRNA1-3’-TCCGGAGACAGCGTTTGGTG-5’, crRNA2-3’-AGGTCCCATCTGGCTCATTG-5’, crRNA3-3’-GTGACTCACCATCCTGACCC-5’. crRNA and tracrRNA were diluted to 100 μM in RNase-free microinjection buffer (10mM of Tris-HCl, pH 7.4, 0.25mM of EDTA). 3 μl crRNA and 3 μl tracrRNA were mixed and annealed in a thermal cycler by heating the mixture to 95°C for 5 minutes and ramped down to 25°C at 5°C/min. Ribonucleic protein (RNP) complexes were prepared by mixing and incubating Cas9 at 0.2 μg/μl with three crRNA/tracrRNA (1μM) in RNase-free microinjection buffer at 37°C for 10 minutes. Embryos were collected from super-ovulated C57BL/6J females mated with C57BL/6J males. Three RNPs were co-delivered into the fertilized eggs by electroporation using NEPA21 super electroporator (Nepa Gene Co., Ltd. Chiba, Japan) under the following conditions: 2 pulses at 40 V for 3 msec with 50 msec interval for poring phase; 2 pulses at 7 V for 50 msec with 50 msec interval for transferring phase. The electroporated embryos were cultured overnight in KSOM medium (EMD Millipore, Billerica, MA) at 37°C and 5% CO2. The next morning, embryos that had reached the two-cell stage were implanted into the oviducts of pseudopregnant ICR CD-1 mothers (Envigo Order code: 030). Pups born were bred with wildtype C57BL/6J mice and were further bred to generate homozygous knockout mice. Homozygous knockout mice were confirmed by stimulating splenocytes with IL-4 and pSTAT6 was measured by flow cytometry.

### Intracellular pSTAT6 staining

To confirm STAT6 KO, mice were euthanized, and spleens were harvested and disrupted mechanically to obtain a splenocyte suspension. Splenocytes were resuspended in RPMI-1640 supplemented with 10% FBS (Sigma-Aldrich Cat# F2442), 1% sodium pyruvate, 1% non-essential amino acids, 1% Penn Strep, 1% L-Glutamine, 2.5% HEPES and stimulated with 50ng/ml IL-4 (R&D Systems Cat: 404-ML) for 15 minutes at 37°C. Splenocytes were then fixed using 1.6% paraformaldehyde for 10 minutes at room temperature. Cells were washed twice with 1x PBS, resuspended in Perm Buffer III (BD Biosciences Cat: 558050) and incubated on ice for 1 hour. Following this, cells were washed three times with FACS buffer (PBS+0.5% BSA+2% FBS+0.1% sodium azide) and stained with the following antibodies for 30 minutes: anti-CD4 PerCP/Cy5.5 (BioLegend Cat: 100540), anti-CD8a Brilliant Violet 510 (BioLegend Cat: 100751), anti-B220 FITC (BioLegend Cat: 103206), anti-STAT6 pY641 Alexa Fluor 647 (BD Biosciences Cat: 558242). Cells were then washed once with FACS buffer and analyzed on Attune NxT flow cytometer. Data were analyzed using FlowJo software.

### Measurement of T and B cell numbers

To measure T and B cell numbers, WT and STAT6 KO mice were euthanized, and spleens were harvested. Spleens were mechanically disrupted to obtain a splenocyte suspension. Splenocytes were stained with eBioscience Fixable Viability Dye eFluor 780 (Thermo Fisher Cat# 65-0865-14) in 1x PBS to exclude dead cells. Splenocytes were washed once and stained with the following antibodies for 30 minutes: anti-CD4 Alexa Fluor 488 (BioLegend Cat: 100529), anti-CD8a Brilliant Violet 510 (BioLegend Cat: 100751), anti-CD19 Brilliant Violet 421 (BioLegend Cat: 115538), anti-CD21 PE/Cy7 (BioLegend Cat: 123419), anti-CD23 PE (BioLegend Cat: 101607). Cells were then washed once with FACS buffer and analyzed on Attune NxT flow cytometer. Data were analyzed using FlowJo software.Murine blood collection and transfusion: Collection and processing of HOD blood was performed as described previously.^30,32^ Details are available in Supporting Information.

### Alum/HEL-OVA vaccination

For immunizations with Alum/HEL-OVA, mice were administered HEL-OVA (100μg) emulsified in aluminum hydroxide (Alhydrogel, InvivoGen Cat# vac-alu-250) via intraperitoneal (i.p.) injection.

### Measurement of anti-HEL antibodies via ELISA

Antibody responses to the Hen egg lysozyme (HEL) portion of the HOD antigen or HEL-OVA were measured by HEL-specific enzyme-linked immunosorbent assay (ELISA), as previously described.^30,32^ Details are available in Supporting Information.

### Statistical Analysis

Graphing of data and statistical analysis were performed using GraphPad Prism software. Mann-Whitney test was done to compare 2 groups. p < 0.05 was considered to be statistically significant and assigned *, whereas p < 0.01, p < 0.001, and p < 0.0001 were assigned **, ***, and ****, respectively.

## Results

### RBC transfusion induces a different IgG class switching profile than vaccination

While previous reports have shown that HOD RBC transfusion leads to production of multiple IgG subclasses in mice,^32^ it is unknown whether the class-switching that is induced by transfused RBCs is similar or different to other class-switch inducing stimuli. We therefore compared antibody production to HOD RBCs with protein emulsified in alum, the gold standard to study IgG isotype class switched antibody responses. Since HOD RBCs express a chimeric protein containing both HEL and OVA, we used HEL-OVA protein emulsified in Alum to minimize potential antigen-specific differences. We measured anti-HEL IgM at 7 days post transfusion/immunization and anti-HEL IgG and subtypes 14 days post transfusion/immunization using endpoint titer ELISAs in order to accurately quantify antibody levels (Figure 1A). Interestingly, transfused RBC consistently led to anti-HEL IgM levels that were at least as high than those observed in response to alum vaccination (Figure 1B). This stood in stark contrast to anti-HEL total IgG levels, which were consistently several logs lower in response to transfusion relative to vaccination (Figure 1C). Both HOD transfusion and Alum/HEL-OVA vaccination led to class switching to each of the four mouse IgG isotypes found in C57BL/6 mice: IgG1, IgG2b, IgG2c and IgG3. However, the relative amounts of antibodies of each subtype differed dramatically between transfusion and vaccination. Relative to vaccination, transfusion with HOD RBCs consistently led to several logs lower anti-HEL IgG1, IgG2b and IgG2c titers (Figure 1D-F). This stood in stark contrast with IgG3 titers, which failed to demonstrate consistently reproducible different titers (Figure 1G). Our data demonstrate that transfusion with HOD RBCs leads to a unique IgG subtype profile in mice that relative to Alum vaccination, favors IgG3 at the expense of IgG1, IgG2b, and IgG2c.

**Figure 1:**
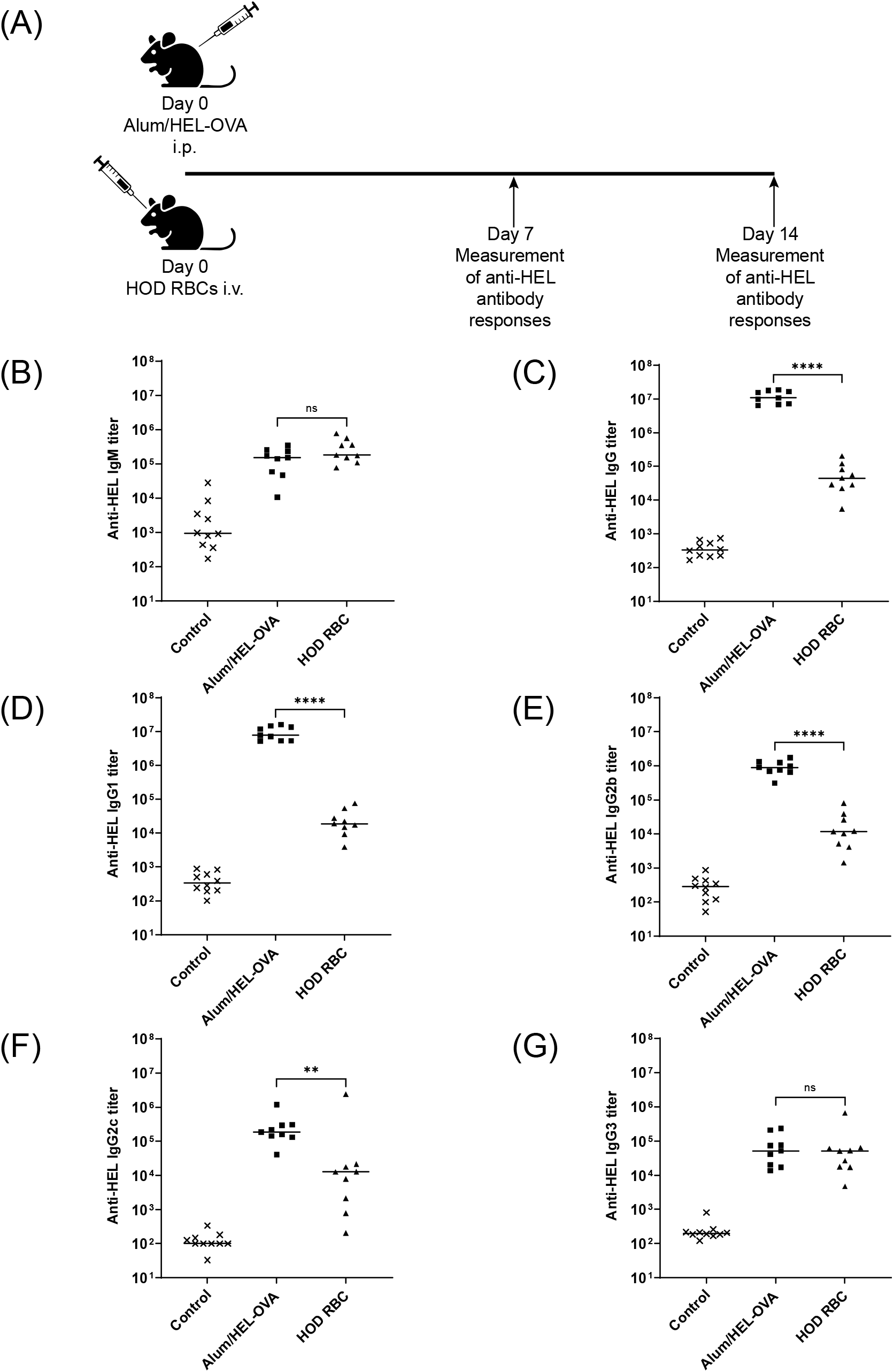
RBC transfusion induces a different IgG class switching profile than vaccination. WT (C57BL/6) mice were either transfused with HOD RBCs or immunized with Alum/HEL-OVA. Anti-HEL titers were measured by ELISA at 14 days post transfusion/immunization. (A) Schematic representation of experimental design (B) Anti-HEL IgM (C) Anti-HEL IgG (D) Anti-HEL IgG1 (E) Anti-HEL IgG2b (F) Anti-HEL IgG2c (G) Anti-HEL IgG3. Each data point represents one mouse. Bars on plots show median values. Figure is representative of 3 independent experiments. *p < 0.05, **p < 0.01, ***p < 0.001, ****p < 0.0001, n.s. p > 0.05.

Given the unique isotype profile observed in response to transfusion, we set out to investigate the mechanisms controlling class switching in this system. STAT6 is a signaling molecule that has been shown to differentially regulate IgG isotype class-switching in mice and humans.^14,33–45,47,48^ We therefore set out to test whether STAT6 signaling similarly regulated class-switching in response to both transfusion and vaccination.

### Generation of STAT6 KO mice

In order to measure the impact of STAT6’s function on class-switching in response to transfusion, we first generated a novel STAT6 KO mouse model using CRISPR/cas9 on a pure C57BL/6 background. Previously generated STAT6 KO mice^34,49^ have used targeting constructs with neomycin resistance cassettes to disrupt STAT6 coding exons.^34,49^ These mice were generated by electroporation into D3 ES cells derived from 129S2/SvPas mouse strain,^34,49^ followed by injection into BALB/c blastocytes and multiple rounds of breeding with C57BL/6 mice to generate STAT6 KO mice on a C57BL/6 background.^49^ This results in a STAT6 deficient mouse that contains not only a neomycin cassette, but also a large swath of tightly linked 129S2 genome surrounding the STAT6 targeted locus. Given the strain specific differences in mice with respect to immune responses^50^ and potential off-target effects of neomycin resistance cassettes,^51^,^52^ we believe that our STAT6 KO mouse model on a pure C57BL/6 background better reflects the wild-type C57BL/6 controls used in our experiments. The CRISPR/cas9 system consists of a crRNA/tracrRNA duplex (gRNA) and cas9 protein, an RNA-guided endonuclease. The crRNA part of the complex binds to targeted exons while the tracrRNA guides the cas9 to the targeted locus. Cas9 endonuclease introduces double-stranded breaks in the targeted region, triggering DNA repair pathways via non-homologous end joining (NHEJ). NHEJ introduces insertions and deletions (indels) and can cause frameshift mutations, leading to disruption of the targeted gene. In order to disrupt STAT6 function, we designed gRNAs to target SH2 coding domains of the STAT6 gene (Figure 2A, Figure 2B). STAT6 SH2 domains are required for STAT6 dimerization and downstream effector function.^53–55^ crRNA/tracrRNA duplexes were complexed with cas9 protein and electroporated into fertilized eggs from C57BL/6 mice, following previously published approaches.^56–58^ Zygotes were transferred into foster mothers and pups were screened for STAT6 mutations and bred to homozygosity (Figure 2A). STAT6 KO homozygous mice were confirmed by stimulating splenocytes with IL-4 and measuring pSTAT6 by flow cytometry within multiple cell populations (Supplementary Figure 1A, Figure 2C). We further characterized the numbers of splenic lymphocyte subsets and observed no significant differences in follicular B cells, marginal zone B cells, CD4+ T cells and CD8+ T cells in STAT6 KO mice compared to WT mice (Figure 2D and Supplementary Figure 1B). Interestingly, while WT and STAT6 KO mice had similar levels of total IgM, STAT6 KO mice showed elevated levels of total IgG (Figure 2E). Our data demonstrate that we have effectively disrupted STAT6 function in our novel STAT6 KO mice.

**Figure 2:**
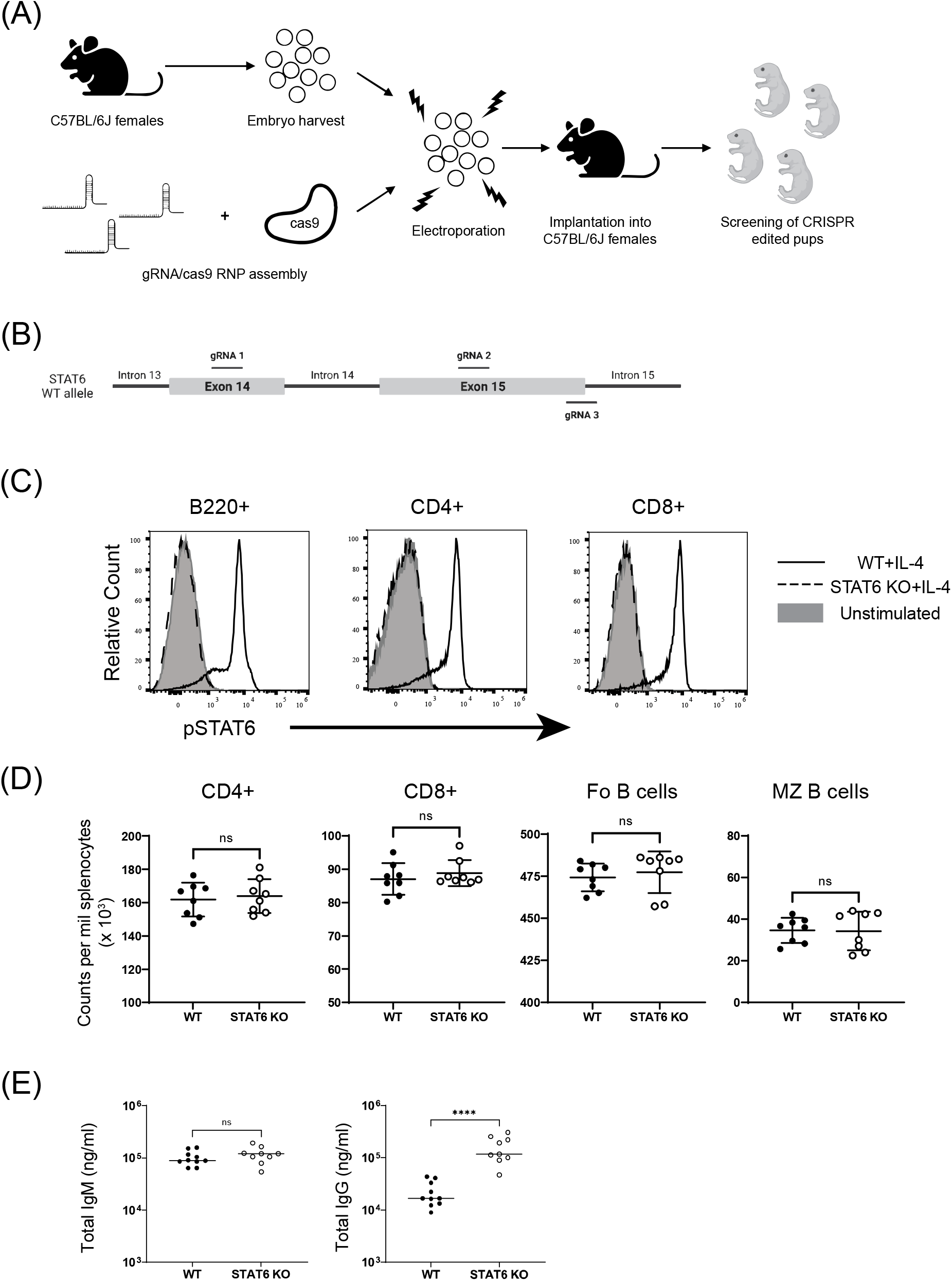
Generation of STAT6 KO mice using CRISPR/cas9 gene editing. (A) A schematic of the KO generation protocol (B) A representative view of the STAT6 targeting strategy showing positions of selected gRNAs (C) pSTAT6 staining measured by flow cytometry following IL-4 stimulation. pSTAT6 in STAT6 KO (dotted line) was compared to WT (solid line). Unstimulated controls are shown using the gray background. (D) Quantification of CD4+ T cells, CD8+ T cells, Follicular B cells and Marginal Zone (MZ) B cells in WT and STAT6 KO spleens. (E) Concentration of total serum IgM and IgG in WT and STAT6 KO mice. *p < 0.05, **p < 0.01, ***p < 0.001, ****p < 0.0001, n.s. p > 0.05.

### STAT6 regulates class switching to all IgG isotypes in Alum/HEL-OVA vaccination

STAT6 is known to be the key transcription factor responsible for Th2 signaling downstream of IL-4 and IL-13. Since protein in alum vaccinations are well known to be Th2 dominant,^14^ we first hypothesized that IgG isotype switching would be altered significantly in STAT6 deficient mice in response Alum/HEL-OVA vaccination. WT and STAT6 KO mice were immunized with Alum/HEL-OVA and anti-HEL IgM and IgG subclass titers were measured (Figure 3A). While total anti-HEL IgG and IgM titers remained similar (Figure 3B and Figure 3C), STAT6 KO mice had significantly lower anti-HEL IgG1 titers and elevated anti-HEL IgG2b and IgG2c titers compared to WT mice (Figure 3D-F), consistent with previous reports.^14^ Interestingly, we found that STAT6 KO mice also had increased anti-HEL IgG3, a novel finding demonstrating that STAT6 also inhibits class switching to IgG3 following Alum/HEL-OVA immunization (Figure 3G).

**Figure 3:**
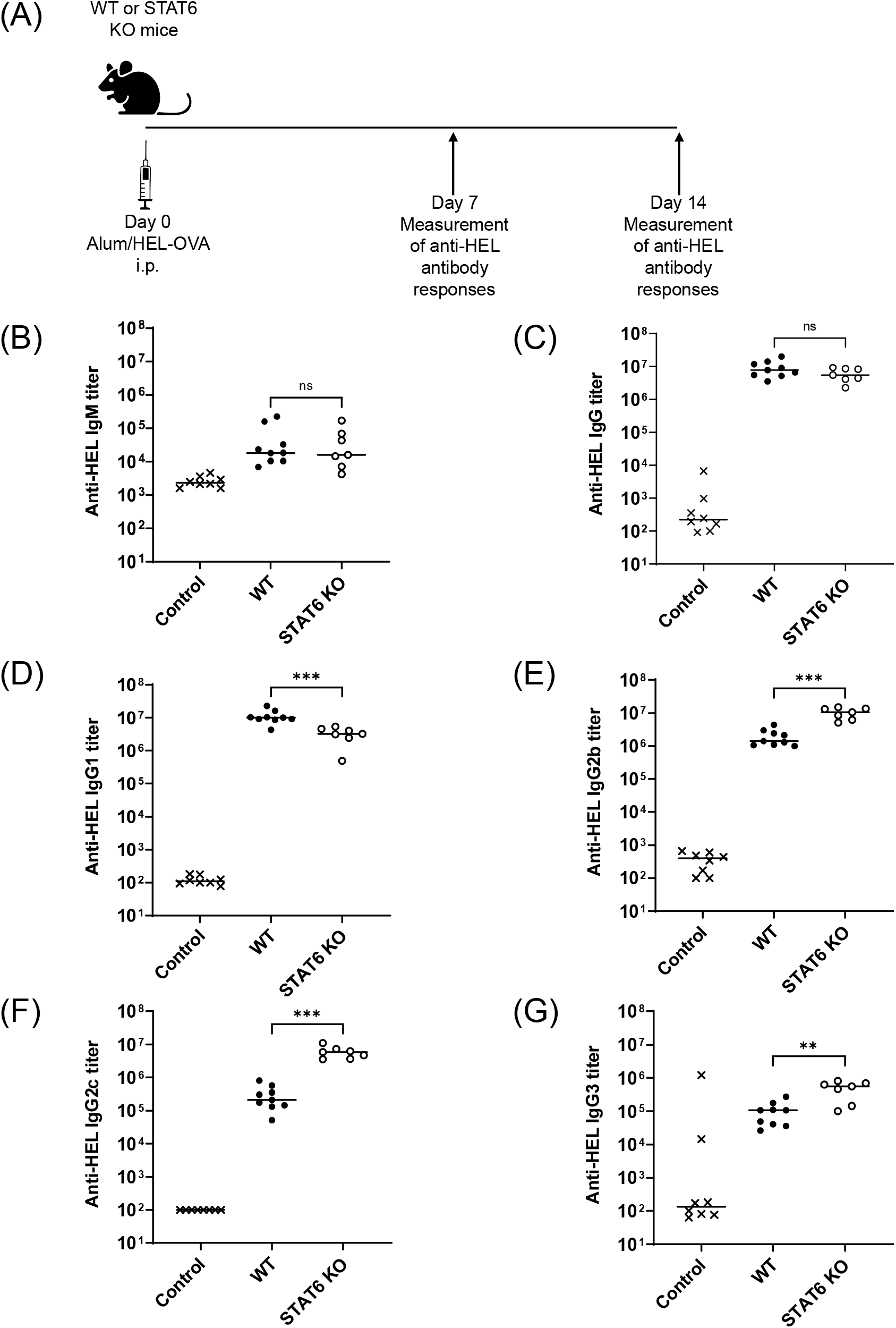
STAT6 regulates class switching in response to Alum/HEL-OVA vaccination. WT and STAT6 KO mice were immunized with Alum/HEL-OVA. Anti-HEL IgG and subtype titers were measured by end point titer ELISAs at 14 days post immunization. (A) Schematic representation of experimental design showing mice immunized with Alum/HEL-OVA (B) Measurement of anti-HEL IgM (C) Measurement of anti-HEL IgG (D) Measurement of anti-HEL IgG1 (E) Measurement of anti-HEL IgG2b (F) Measurement of anti-HEL IgG2c (G) Measurement of anti-HEL IgG3. Each data point represents one mouse. Bars on plots show median values. Figure is representative of 3 independent experiments. *p < 0.05, **p < 0.01, ***p < 0.001, ****p < 0.0001, n.s. p > 0.05.

### STAT6 has a minimal impact on class switching to most IgG isotypes in response to HOD transfusion

In order to determine the impact of STAT6 on transfusion-induced class switching, WT and STAT6 KO mice were transfused with HOD RBCs and anti-HEL antigen specific titers were measured. We found that WT and STAT6 KO mice had similar anti-HEL IgM and IgG titers (Figure 4B and Figure 4C). We then measured anti-HEL IgG subclass titers 14 days post transfusion. To our surprise, STAT6 deficient mice had similar levels of IgG1, IgG2c and IgG3 in response to HOD transfusion (Figure 4D, Figure 4F and Figure 4G). We also observed that STAT6 KO mice showed increased IgG2b (Figure 4E). Collectively, our data demonstrates that transfusion-induced IgG class switching occurs in a largely STAT6-independent manner, suggesting that alternative pathways that are largely STAT6-independent regulate IgG class switching in response to RBC transfusions.

**Figure 4:**
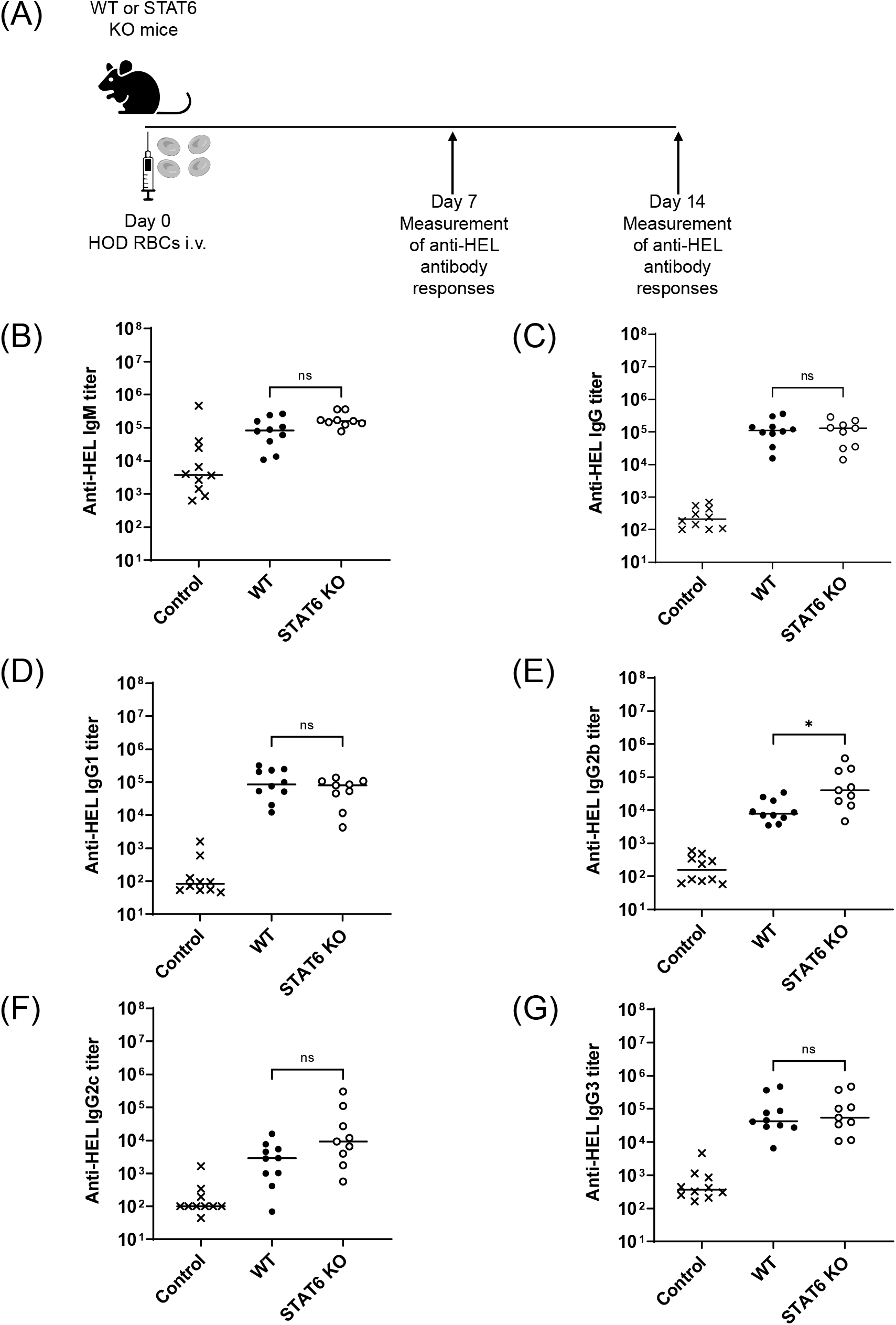
STAT6 is mostly not required for class switching in response to HOD RBC transfusion. WT and STAT6 KO mice were transfused with HOD RBCs. Anti-HEL IgG and subtype titers were measured by end point titer ELISAs at 14 days post transfusion. (A) Schematic of experimental design showing mice transfused with HOD RBCs (B) Measurement of anti-HEL IgM (C) Measurement of anti-HEL IgG (D) Measurement of anti-HEL IgG1 (E) Measurement of anti-HEL IgG2b (F) Measurement of anti-HEL IgG2c (G) Measurement of anti-HEL IgG3. Each data point represents one mouse. Bars on plots show median values. Figure is representative of 3 independent experiments. *p < 0.05, **p < 0.01, ***p < 0.001, ****p < 0.0001, n.s. p > 0.05.

## Discussion

Our data in the HOD mouse model demonstrate that transfusion of HOD RBCs leads to consistent generation of IgG. However, we show that, compared to Alum/HEL-OVA vaccination, HOD RBC transfusion induces similar IgG3 titers but dramatically lower IgG1, IgG2b and IgG2c titers. This demonstrates that even though all IgG isotypes are produced in response to transfusion, the levels of a given isotype generated in response to transfused RBCs are different than those generated in response to protein in Alum vaccination.

Given the observed differences in isotype class switching induced by transfusion vs. vaccination, we were interested in determining whether the molecular regulators of class switching might also be different between these immune stimuli. Since STAT6 is a well-known regulator of IgG1 and IgG2 subtype class switching in multiple different vaccination and infectious settings^14,47,48,33–45^, we created a novel STAT6 deficient mouse model to directly test whether STAT6 played a similar role in regulating isotype class-switching in response to transfusion. We used CRISPR/cas9 to target STAT6 in C57BL/6J embryos and confirmed successful disruption of STAT6 function. These mice represent, to our knowledge, the first STAT6 deficient strain generated on a pure C57BL/6 background. STAT6 deficient mice had similar levels of baseline polyclonal IgM, but elevated levels of baseline polyclonal IgG. This suggests that in response to the normal microflora found in our specific pathogen free (SPF) colony at University of Virginia, STAT6 deficient mice tend to have higher circulating total IgG levels relative to their wild type counterparts.

Having established the baseline polyclonal antibody production in STAT6 KO mice, we next asked what the impact of STAT6 deficiency in the production of antigen-specific antibody production in response Alum/HEL-OVA vaccination. Consistent with previous publications^14^, STAT6 KO mice that were vaccinated with Alum/HEL-OVA expressed similar levels of antigen-specific IgM and antigen-specific total IgG. Furthermore, STAT6 KO mice expressed lower levels of antigen-specific IgG1 while expressing significantly elevated levels of antigen-specific IgG2b, IgG2c and IgG3 subclasses in response to Alum vaccination. These data demonstrate that STAT6 plays a significant role in the relative class switching to all IgG isotypes in response to Alum vaccination, supporting IgG1 antigen-specific antibody production while suppressing IgG2b, IgG2c and IgG3 antigen-specific antibody production.

Importantly, STAT6 deficiency had a much different impact on antigen-specific IgG subclass production in response to transfused RBCs. Unlike what was observed in response to Alum vaccination; anti-RBC IgG1, IgG2c and IgG3 levels were not consistently or robustly different between wild type mice and STAT6 KO mice. While anti-RBC IgG2c levels did trend somewhat lower, they failed to consistently show a significant difference. We did however observe a small but reproducible enhancement in IgG2b levels in absence of STAT6. Thus, our data demonstrate that STAT6 clearly suppressed IgG2b (and to some degree IgG2c) class-switching in response to transfusion. However, there was no significant impact on anti-RBC IgG1 and IgG3 production in STAT6 deficient mice. This demonstrates that the molecular regulation of the antigen-specific IgG1 and IgG3 class-switching in response to transfusion is quite different from that induced by Alum vaccination.

There are several potential mechanisms that might account for the differences that we observed between HEL/OVA in Alum vaccination and HOD RBC transfusion. One potential explanation for the differences we observe may be associated with differences in antigen doses between HEL/OVA in Alum vaccination and HOD transfusion. Although certainly a possibility, we do not favor antigen dose differences as a major explanation for our results for several reasons. First, experiments in our lab using a range of varying doses of HOD RBCs showed no significant drop-off in antibody titers for the doses tested (data not shown). Second, significant variation in the dose of Alum/HEL-OVA had only a small effect on overall titers and did not impact isotype distribution (data not shown). Thus, we do not think that intrinsic differences in antigen dosing account for the differences in class switching observed.

Another potential difference that might account for our observed differences in class-switching is the different route of antigen exposure: with the vaccination given intra-peritoneally (i.p.) while the transfusion is given intravenously (i.v.). Importantly, direct comparisons of Alum and transfusion are not possible since Alum cannot be given i.v. in mice due to toxicity, and i.p. transfusion of HOD RBCs fails to generate any measurable anti-RBC alloantibodies (data not shown). However, we do not think that the route of exposure accounts for the differences we observed since Alum used via other routes (intra-muscularly) gives similar results to i.p., and i.v. protein vaccination with i.v.-compatible adjuvants such as the Sigma Adjuvant System show similar robust induction to IgG1 and IgG2 subclasses relative to IgG3 to what we have observed in the i.p. Alum setting (data not shown). Furthermore, STAT6 controls IgG1 antigen-specific class switching to multiple different infectious agents that infect multiple different sites, including intestinal helminth infection,^48,59^ intranasal pox viral vaccination,^45^ and upper respiratory flu viral infection.^60^ Thus we believe that the i.p. vaccination gives similar results to vaccination and infection at many other sites. However, it is certainly possible that i.v. exposure to antigens that occurs via transfusion may account for some of the observed differences in class switching. Importantly, RBC transfusions, by definition entail i.v. antigen exposure via RBCs, and thus our model accurately depicts the unique properties of transfusion-associated induction of class-switching.

Though route of immunization and antigen dosing are potential explanations for the differences in IgG class-switching we have observed, we favor the hypothesis that antigens presented on transfused RBCs represent a unique costimulatory signal to the immune system. This is particularly true when sterile RBCs are transfused in the absence of exogenous adjuvants as is the case for our studies herein and for most clinical transfusions. We have used HEL/OVA in Alum vaccination as a comparator in our studies because protein in Alum vaccination has long served as the gold standard for the scientific study of class-switching, and protein in Alum vaccination continues to be the vaccine approach used in most childhood vaccines. Importantly, HEL/OVA in Alum vaccination faithfully represents the vast majority of the published data looking at infectious and vaccine induced class-switching in so far as STAT6 clearly supporting IgG1 class-switching and suppressing IgG2 isotype class switching. The key point here that we would like to emphasize is not that the IgG class-switching induced transfusion is different from the IgG class-switching induced by HEL/OVA in Alum per se, but rather that the class-switching induced by transfusion is different from virtually every other infectious and adjuvant-based system that we have looked at and that has been studied (Alum included). Ultimately, we hypothesize that foreign antigens that are expressed on transfused RBCs stimulate the immune system in a unique manner relative to most other vaccination or infectious stimuli, driving a relatively robust IgG3 response but dramatically weaker IgG1, IgG2b and IgG2c responses. Our data herein demonstrate that STAT6 fails to play a significant role in the production of most of the class-switched IgG antibodies generated in response to HOD transfusion, and certainly not IgG3 and IgG1. Given the unique nature of the IgG class switching observed in response to transfusion, we are particularly interested in future experiments aimed at understanding both the cellular and molecular regulators of IgG production in response to RBC-expressed foreign antigens.

## Acknowledgements

These studies were supported by grants to C.J.L from National Institutes of Health (R01 HL134691) and to C.J.L and J.E.H (P01 HL132819). A.P. was supported in part through the Immunology Training Grant at UVA from the National Institutes of Health (5T32AI007496).

The authors would also like to thank the UVA Genetically Engineered Murine Model (GEMM) core for their support.

The authors would also like to thank Mrs. Jennifer Taylor, Mr. Asher Compton and the animal care staff at the University of Virginia School of Medicine.

## Notes

**<u>Conflicts of Interest:</u>** The authors report no conflicts of interest

### Competing Interest Statement

The authors have declared no competing interest.

