## Supporting Information for "Class switching is differentially regulated in RBC alloimmunization and vaccination"

**Materials and Methods:**

Measurement of anti-HEL antibodies via ELISA: High-binding polystyrene plates (Corning # 9018) were coated for 1 hour at 37°C with 10 μg/ml HEL (Sigma-Aldrich Cat: L6876) in PBS. Plates were then washed (0.05% Tween-20 in PBS) and incubated with blocking buffer (2% BSA and 0.05% Tween-20 in PBS) overnight at 4°C. Sera samples were serially diluted (4-fold dilutions starting at 1:50, diluted 12 times) in blocking buffer and incubated in coated plates for 1 hour at room temperature. Wells were then incubated for 1 hour at room temperature with one of the following horseradish peroxidase conjugated secondary antibodies: goat anti-mouse IgM, goat anti-mouse IgG, goat anti-mouse IgG1, goat anti-mouse IgG2b, goat anti-mouse IgG2c, goat anti-mouse IgG3 (Jackson ImmunoResearch Codes: 115-035-075, Jackson ImmunoResearch Codes: 115-035-008, Jackson ImmunoResearch Codes: 115-035-205, 115-035-207, 115-035-208, 115-035-209 respectively). Wells were developed using 3,3′,5,5′-Tetramethylbenzidine (TMB) substrate (SeraCare Cat# 52-00-03) and quenched with 2 N $H_2SO_4$ after 10 min. Optical densities were measured at 450 nm. End-point titers were calculated using GraphPad Prism through interpolation of the cutoff value from the fit of the optical density versus (1/serum dilution) curve for each sample using the "plateau followed by one-phase decay" model. The cutoff value was defined as the average plus 3 standard deviations (SDs) of signals from background wells (i.e., signal values from wells incubated with blocking buffer alone). O.D values that started off below background and unable to be interpolated were assigned a titer value of 100.

Measurement of total serum IgG and IgM: In order to measure total serum IgM and IgG levels in WT and STAT6 KO mice, serum was collected from naïve 8-week-old mice. Total IgM was measured using IgM mouse uncoated ELISA kit (ThermoFisher Cat: 88-50470-22) and total IgG was measured using IgG (Total) mouse uncoated ELISA kit (ThermoFisher Cat: 88-50400-22). ELISA was performed based on manufacturer's instructions.

Murine blood collection and transfusion: Blood from HOD mice was aseptically collected by cardiac puncture into the anticoagulant citrate phosphate dextrose adenine (CPDA-1, Boston Bioproducts IBB-420). The final volume was adjusted to 20% CPDA-1 (v/v). Collected HOD blood was leukoreduced using whole blood cell leukoreduction filter (Pall, AP-4851). Leukoreduced blood was centrifuged at 1200 x g for 10 minutes, adjusted to a final hematocrit of 75% and stored 4°C for 12 days. Recipient mice received 100μl of 75% Hct HOD RBCs intravenously via retroorbital injection.

**Supplementary Figure 1**

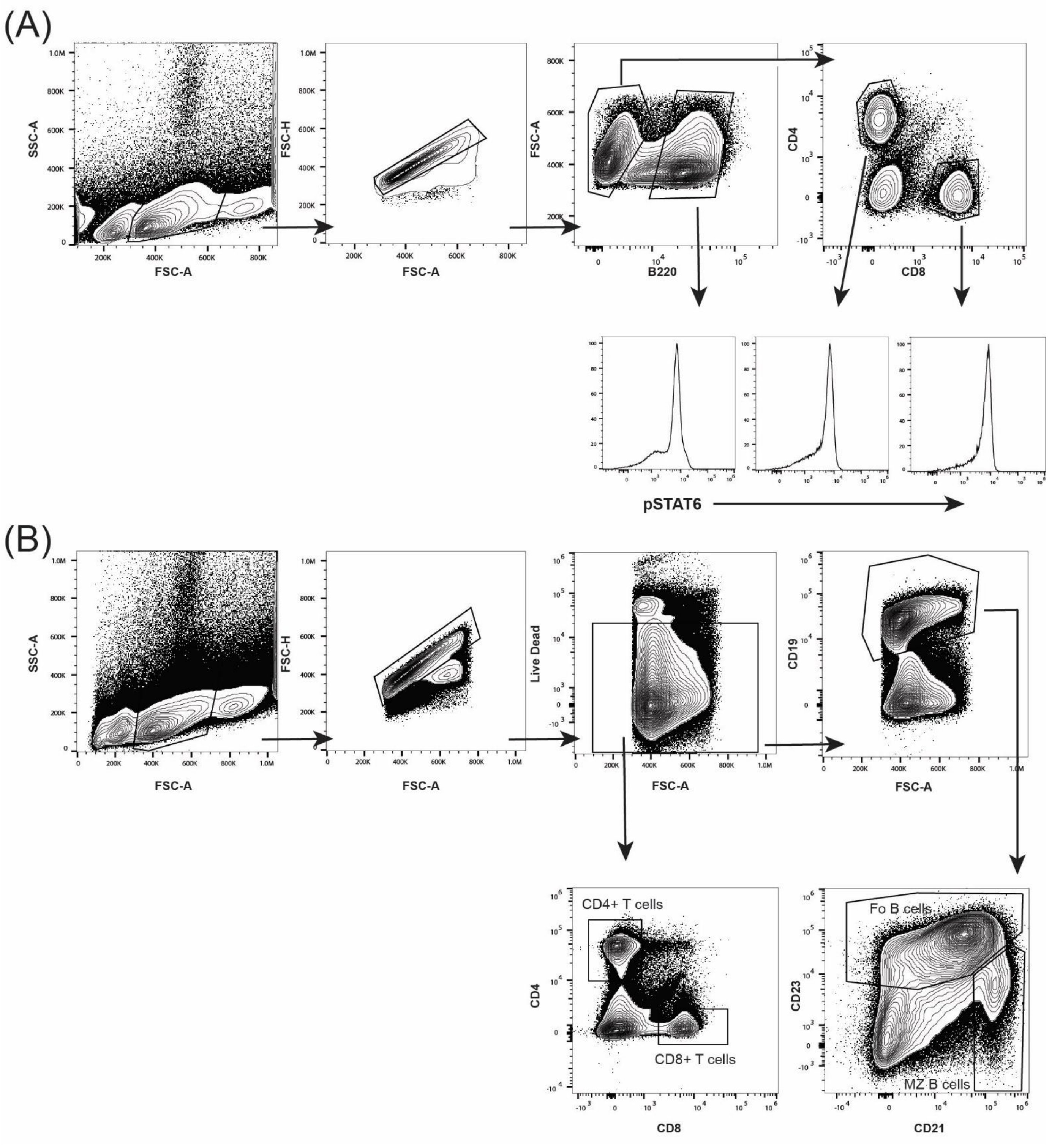

**Supplementary Figure 1:** (A) Gating strategy for measurement of pSTAT6 by flow cytometry. pSTAT6 levels were measured by flow cytometry in response to IL-4 stimulation in CD4+ T cells, B220 + B cells and CD8+ T cells from WT and STAT6 KO mice. (A) Gating strategy for detection of pSTAT6 within specific cell types. Representative figure shows gating strategy using WT mice. (B) Gating strategy for measurement of CD4+ T cells, CD8+ T cells, follicular B cells and marginal zone B cells.
